# A novel supergene controls queen size and colony social organization in the ant *Myrmica ruginodis*

**DOI:** 10.1101/2025.03.24.644106

**Authors:** Hanna Sigeman, Perttu Seppä, Philip A. Downing, Matthew Webster, Heikki Helanterä, Lumi Viljakainen

## Abstract

In some ant species, the size of queens is dimorphic, with larger queens dispersing to form single-queen colonies and smaller queens remaining in natal colonies as part of multiple-queen societies. These alternative reproductive strategies influence colony kin structure and worker behavior, and their many independent origins make them ideal systems to study parallel evolution of genome organization and adaptability. We investigate the genetic basis of alternative reproductive strategies in the queen-size dimorphic ant *Myrmica ruginodis* by sampling 95 queens from 31 colonies in southern Finland. Whole-genome sequencing revealed a novel 9 Mb supergene associated with both queen size and social organization. Queens homozygous for the AA haplotype were larger and found only in single-queen colonies, while queens in multiple-queen colonies were smaller and contained only AB and BB genotypes. This supergene is not homologous to previously identified supergenes in ants, suggesting it was formed by a unique evolutionary pathway.

## Introduction

Alternative reproductive strategies are common across many species, involving traits such as size differences, behavioral tactics like sneaker males, or variations in sex allocation and parental care (Oliveira et al., 2008). In ants, these strategies may manifest as discrete queen phenotypes living in colonies with different social organizations. The ancestral reproductive strategy involves a queen morph capable of dispersal and independent colony founding (Hölldobler & Wilson, 1990). From this (*monogynous*) single-queen colony structure, queens have repeatedly evolved into philopatric, and often smaller, morphs living in (*polygynous*) multi-queen colonies. These polygynous queens often forego the risky life stage of independent dispersal, and instead become one of several reproductive queens in their natal colony (e.g. Heinze & Tsuji, 1995; Rüppell & Heinze, 1999; Wolf & Seppä, 2016). These two strategies have gained considerable interest from researchers, partly because the number of queens influences genetic relatedness within colonies, shaping worker behavior to optimize their inclusive fitness (e.g. Crozier & Pamilo, 1996). Additionally, these traits offer a striking example of parallel evolution, providing a framework for testing hypotheses on genome organization and adaptive flexibility. While the genetic mechanisms underlying colony social structure have been elucidated in five ant lineages—each involving large supergenes that govern the traits associated with the two queen morphs (Braim, 2015; Brelsford et al., 2020; Errbii et al., 2024; Lagunas-Robles et al., 2021; Lajmi et al., 2024; Purcell et al., 2014, 2014; Scarparo et al., 2023; J. Wang et al., 2013) —many other lineages with multiple queens and/or queen-size dimorphism remain unexplored. Investigating additional systems will broaden our understanding of genome dynamics, adaptability, and parallel evolutionary processes.

The common Palearctic red ant *Myrmica ruginodis* (Nylander) is characterized by queen-size dimorphism, which is linked to distinct reproductive strategies. Large *macrogyne* queens disperse from their natal colonies to join mating swarms and establish new colonies independently, where they are the sole reproductive female in monogynous colonies. In contrast, smaller *microgyne* queens are found in polygynous colonies, where multiple queens coexist and reproduce. Microgynes tend to mate within or near their natal colonies to which they are often recruited back as reproductive queens, and new colonies are founded by splitting old ones (*budding*, Brian & Brian, 1949, 1955). The association between queen morph and colony social organization in *M. ruginodis* is not absolute, however. Although queen-size distribution is bimodal and queens are on average larger in monogynous than in polygynous colonies, size distributions are overlapping, and intermediate queens cannot always be classified to either morph with certainty. Moreover, while most macrogynes disperse and most microgynes are recruited back to their natal colony, queens of both types sometimes deviate from these patterns (Elmes, 1991; Wolf et al., 2018).

Although the genetic mechanisms underlying queen-size and colony social organization in *M. ruginodis* are currently unknown, previous genetic studies provide important background information to the system. Allozymes and DNA microsatellite data have revealed only weak genetic differentiation between monogynous and polygynous colonies (Wolf, 2016), explained partly by males from both colony types joining mating swarms (Wolf & Seppä, 2016). Most monogynous colonies are simple families, where workers are daughters of the resident queen. However, sometimes the resident queen is not the mother of all workers (Seppä, 1994). Multiple matrilines within monogynous ant societies have also been found in other ant species, and can be explained by worker recruitment of a new queen in colonies where the previous queen has died (so-called *serial polygyny*; reviewed in Heinze & Keller, 2000). High relatedness among queens in polygynous colonies confirms that most queens rejoin their natal colony after mating, but unrelated queens also occur in them (Seppä, 1994; Walin & Seppä, 2001; Wolf et al., 2018). Lastly, queen-size in *M. ruginodis* is a heritable trait and thus expected to be fully or partially under genetic control (Wolf, 2016).

Here we identify a novel 9 Mb supergene in *M. ruginodis* and show that it controls both queen-size and colony social organization. The supergene is non-homologous to previously identified and well-characterised supergenes controlling monogyny/polygyny in ants (*Cataglyphis* (Lajmi et al., 2024), *Formica* (Purcell et al., 2014), *Pogonomyrmex* (Errbii et al., 2024), and *Solenopsis* (J. Wang et al., 2013)), suggesting an independent origin within the *Myrmica* lineage. It is also non-homologous to a second supergene in *Formica*, so far detected only in *F. cinerea*, which control queen-size within polygynous colonies (Scarparo et al., 2023). Our discovery solves the long-standing mystery of alternative reproductive strategies in *M. ruginodis* and introduces a social supergene in a new ant lineage, adding to the five already known.

## Methods

### Nomenclature

In this paper, we follow the British tradition of referring to wingless adult reproductive females as *queens*, and newly hatched and winged ones as *gynes*. Moreover, with *single-queen* and *multiple-queen* colonies we refer only to the observed number of queens. Fast queen turnover, colony-founding by budding with concurrent queen dispersal, and failure to sample all queens may all lead to a situation where the observed queen number does not correspond to the true number of breeding queens in the colony (see above). We therefore distinguish further and call colonies where all workers are offspring of a single queen as *monogynous* and those with multiple reproducing queens as *polygynous*.

## 1. Whole-genome sequencing data: sampling and analyses

### 1.1 Field work, morphometrics, DNA Extraction, and WGS sequencing

In 2023 we collected *M. ruginodis* queens and gynes from seven colonies from the well-studied TV/Leimann population in the vicinity of Tvärminne Zoological station, Hanko, SW-Finland (Table S1). The Hanko area is characterized by dry pine forest, where *M. ruginodis* workers build their colonies in a thin layer of vegetation above the sandy bottom. This allows excavation of entire colonies, a method that rarely lets queens escape collection (Seppä, 1994; Wolf et al., 2018). We manually sorted each colony in the lab and collected all queens and gynes, as well as a subset of workers. We decapitated each queen and gyne with a sterilized scalpel and photographed the head from a dorsal view using a microscope camera. We used a cell counting chamber (a glass plate with a 0.02 mm grid) as a scale and extracted precise measurements of the head width (maximum width directly above the eyes) using the software ImageJ2 (Rueden et al., 2017) (Table S1). A cutoff value of 1.06 mm has been used previously to distinguish macrogynes and microgynes (Elmes, 1991; Wolf et al., 2018). As many of the measured individuals were close to this, we here categorized queens and gynes >1.08 mm as macrogynes, <1.04 mm as microgynes and 1.04–1.08 mm as intermediates. The bodies of the queens/gynes, and the whole workers were frozen in liquid nitrogen and stored in -80°C. DNA from 11 queens and 6 gynes was extracted using DNeasy Blood & Tissue Kit (QIAGEN), and whole-genome sequenced on an Illumina NovaSeq X Plus (PE150) platform with a Plant and animal whole genome library preparation (350bp). We also whole-genome sequenced 18 *M. ruginodis* workers, each collected from a different colony in the TV/Leimann population (Table S1), using the same DNA extraction and sequencing protocol.

### 1.2 Chromosome-level pseudo-assembly of the M. rubra reference genome

The genome of *M. rubra*, a sister-species to *M. ruginodis*, was recently assembled as part of the Global Ant Genomics Alliance (GAGA) consortium initiative to generate reference genomes for lineage-representatives of each ant genus (Boomsma et al., 2017). To increase contiguity of this scaffold-level assembly (N50: 1.4 Mb; 477.0 Mb; Table S2), we corrected and scaffolded the genome sequences using the chromosome-level reference genome of *Solenopsis invicta* (N50: 26.2 Mb, total length: 378.1 Mb; Table S2). The two species have an estimated divergence time of ∼100 Mya (Prebus & Rabeling, 2024). Specifically, the PacBio reads underlying the *M. rubra* reference genome were transformed to FASTQ format using bam2fastq from the PacBio BAM toolkit v3.0.0 (https://github.com/PacificBiosciences/pbtk), and corrected using canu v2.2 (Koren et al., 2017). The corrected reads had 27.09 times sequence depth (Table S3). We used ragtag correct v2.1.0 (Alonge et al., 2022) to break scaffolds in the *M. rubra* genome at potential misassembly sites using *S. invicta* as a target genome. To avoid breaking scaffolds in incorrect places and within genic regions, we provided the corrected PacBio reads as supporting evidence as well as the *M. rubra* gene annotation file (GFF). The corrected reference genome was then scaffolded using *S. invicta* as a target genome. The scaffolded *M. rubra* reference genome had an N50 value of 18.3 Mb and a total length of 477.1 Mb (Table S2). The genome coordinates of the *M. rubra* gene annotation were translated to the new genomes using ragtag updategff, first from the original to the corrected version, and then from the corrected version to the scaffolded version. Genome statistics were calculated with QUAST v5.2.0 (Gurevich et al., 2013). We assessed the genome completeness of the original and scaffolded *M. rubra* genome using BUSCO v5.4.7 (Simão et al., 2015) using the hymenoptera_odb10 dataset (eukaryota, 2024-01-08), and found that the scaffolding did not lower the quality of the reference genome (original *M. rubra*: 94.1% complete BUSCOs; *M. rubra* scaffolded: 94.2% complete BUSCOs; Table S4).

### 1.3 Alignment and variant calling

Whole-genome sequencing data from the 17 *M. ruginodis* queens/gynes and 18 *M. ruginodis* workers (Table S1) were aligned to the scaffolded *M. rubra* reference genome using NextGenMap v0.5.5 (Sedlazeck et al., 2013), sorted with samtools v1.16.1 (Danecek et al., 2021), and deduplicated with picard v2.27.5 (http://broadinstitute.github.io/picard/). We calculated alignment statistics based on the deduplicated BAM files using samtools stats. The sequencing depth for the queens/gynes varied between 16.1x and 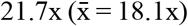, and for the workers between 11.2x and 13.0x (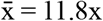Table S5). Variants were called for each sample separately using deepvariant v1.4.0 (Yun et al., 2021), followed by joint variant calling with glnexus v1.4.1 (Yun et al., 2021). We normalized the VCF file with bcftools norm v1.16 (Danecek et al., 2021) and filtered using vcftools v0.1.17 (--min-alleles 2 --max-alleles 2 --minQ 20 --minDP 3 --max-missing 0.6 --remove-filtered-all) (Danecek et al., 2011). We calculated pairwise F_ST_ between macrogynes, microgynes, and intermediates, across 10kb windows with vcftools.

To visualize genotypes across the supergene region on chromosome 5, we made a rolling window PCA plot by first filtering the original VCF file for this chromosome, and then splitting the new VCF file into separate files for each 50 kb window. These 50 kb VCF files were transformed to PLINK format using plink v2.00a6 (www.cog-genomics.org/plink/2.0/) (Chang et al., 2015), followed by a PCA analysis using plink for each window. PCA axis 1 was plotted along the chromosome using ggplot2 v3.5.1 (Villanueva & Chen, 2019) in R v4.4.1 (R Core Team, 2022). Heterozygosity across chromosome 5 was calculated with vcftools using the --het option (Table S6). Linkage disequilibrium between SNPs is expected to be constant throughout supergene regions, while rapidly decaying with distance in normally recombining (non-supergene) regions. We therefore calculated decay of linkage disequilibrium for (i) each chromosome (1-16), for the (ii) supergene and (iii) non-supergene regions of chromosome 5 using PopLDdecay v3.43 (Zhang et al., 2019). For each resulting file, we used the script Plot_OnePop.pl from the same program to calculate r^2^ values across bins using the default settings. We used values from all chromosomes except chromosome 5 to calculate genome-wide r^2^ values.

### 1.4 Supergene haplotype gene content

All whole-genome sequenced samples (n = 17 queens/gynes and n = 18 workers; Table S1) were categorized as AA, AB, or BB genotypes according to which SNP genotypes were most common within the supergene region (see Results). Fixed haplotype-specific within-exon SNPs (heterozygous in AB individuals, while having opposing homozygosity between all AA and BB individuals) were extracted. First, all SNPs that were heterozygotic in all AB individuals were extracted using snpSift v5.1d (Cingolani et al., 2012). Second, we calculated F_ST_ between the AA and BB individuals using vcftools with the –weir-fst-pop option and selected only SNPs with F_ST_ values of 1. Lastly, we extracted the longest transcript from each gene in the GFF file using the AGAT v1.0.0 (Dainat, n.d.) toolkit and the script agat_sp_keep_longest_isoform.pl, and extracted only the SNPs within CDS regions (n = 594 SNPs). Then we used orthofinder v2.5.5 (Poplin et al., 2018) to find orthologs between selected ant species with high-quality reference genomes and their associated gene annotations (*Atta, Solenopsis, Pogonomyrmex, Myrmica*; Table S2).

### 1.5 Synteny

We analyzed synteny between four lineages where the genome region for supergenes underlying colony social organization is known: *Formica* (Brelsford et al., 2020), *Myrmica* (this study), *Pogonomyrmex* (Errbii et al., 2024), and *Solenopsis* (Yan et al., 2020) (see Table S2, S7 for genome and annotation versions, as well as supergene genome regions for each reference genome). We used the GENESPACE v1.3.1 (Lovell et al., 2022) analytical pipeline, which relies on orthofinder to identify orthologs, and MCScanX (Y. Wang et al., 2012) (downloaded from GitHub 26^th^ of September 2024) for visualization of syntenic blocks. Prior to analysis with GENESPACE we filtered each GFF file for the longest transcript for each gene using the script agat_sp_keep_longest_isoform.pl from the AGAT toolkit. The output from GENESPACE was plotted using Circos v0.69-8 (Krzywinski et al., 2009).

## 2. Confirming association between queen morph and supergene with targeted genetic markers

### 2.1 Field work and morphometrics

To increase our sample size of *M. ruginodis* queens we returned to the same area in SW-Finland in 2024 to collect additional colonies. In total, we collected 17 colonies from the TV/Leimann population and 7 colonies from the nearby Täktom population (populations separated by >10 km; Table S1). All queens were stored in 96% ethanol until decapitation and morphometric photography (following the same protocol as above), and then stored in -20°C along with a subset of workers from each colony.

### 2.2 Supergene genotyping through PCR experiments

Using the 17 whole-genome sequenced *M. ruginodis* queens/gynes, we designed diagnostic PCR primers for the supergene, capable of distinguishing between AA, AB, and BB genotypes through a B-specific 2 bp insertion (primer pair “IL16”; Table S8). The same individuals were used to confirm the DNA product accuracy through PCR and Sanger sequencing. The resulting electropherogram files (ab1 format) were phased using the R package sangerseqR v1.40.0 (Hill et al., 2014), using one of the whole-genome sequenced queens with an AA supergene genotype (MrugQ14; Table S1) as a reference. The sequences were trimmed to the region between 50 bp and 200 bp and aligned using muscle v5.1 (Edgar, 2022). The region around the diagnostic 2 bp indel was plotted for all samples using the R package ggmsa v1.10.0 (Zhou et al., 2022), which showed a perfect correlation to the supergene genotype assignment from the whole-genome sequencing data (Table S1; Figure S1). We then extracted DNA from all remaining queens (n = 78; Methods section 2.1; Table S1) using DNeasy Blood & Tissue Kit (QIAGEN), and used the (1) IL16 primers to genotype them for the supergene and (2) the universal mitochondrial primer pair LCO1490 and HCO2198 (Folmer et al., 1994) to confirm the species identity. Details on primer pairs and PCR protocol are provided in Table S8. For the mitochondrial marker, targeting the COI gene, we aligned the sequences using muscle and then trimmed the aligned sequences to the region between 22bp and 680bp. We used the R package ape v5.8 (Paradis et al., 2004) to calculate pairwise sequence distance using the default K80 method between all individuals. The sequence distances were very low between all individuals (ranging between 0 and 0.006), confirming that they are all *M. ruginodis* (Figure S2).

### 2.3 DNA microsatellite genotyping

To confirm the kin structure in single-queen colonies (see above), we investigated the genetic relationship between workers using five microsatellite markers (Table S9). All markers were originally designed for other *Myrmica* species (Azuma et al., 2005; Henrich et al., 2003; Zeisset et al., 2005) and also tested directly in *M. ruginodis* (Wolf et al., 2018). DNA was extracted using 6% Chelex and Proteinase K. For details on the PCR protocol for each marker see Wolf et al., (2018). We analyzed 10-11 workers from thirteen single-queen colonies, and 3-8 workers from five single-queen colonies (Table S10). As a control, we also investigated worker relationships in four multiple-queen colonies (Table S10). Genotypes were predicted for each marker with the GUI program STRyper v1.2.2 (Peccoud, 2023). Each peak was manually inspected in STRyper, and incorrect genotype calls were adjusted (Tables S11-S15 show the original and adjusted genotype calls for each marker; Table S10 shows the final scored genotypes). We tested the genetic relationship among workers by running all 199 samples simultaneously in Colony v2.0.7.1 (J. Wang, 2018). We used the default settings for haplodiploid organisms in the analysis and an allelic dropout rate and false allele rate of 5%. As a small proportion of *M. ruginodis* queens mate multiple times (∼16%; Seppä, 1994), we ran the algorithm both with and without polyandry (i.e. maternal polygamy).

## Results and Discussion

### Association between queen size and novel supergene polymorphism

To identify genome regions associated with queen size, we aligned genome sequences from the 17 *M. ruginodis* queens/gynes to the pseudo-scaffolded chromosome-level *M. rubra* reference genome. The head sizes ranged between 0.92 and 1.14 mm, thereby categorizing 3 as macrogynes, 4 as intermediates, and 10 as microgynes (Table S1). We searched for local peaks of differentiation between these three queen types across all chromosomes using F_ST_, which revealed a highly differentiated region on chromosome 5 spanning 9 Mb (20.2–29.2 Mb; Figure 1). This region was significantly differentiated between macrogynes and microgynes (Figure 1B), somewhat differentiated between macrogynes and intermediates (Figure 1A), but not between intermediates and microgynes (Figure 1C).

**Figure 1.**
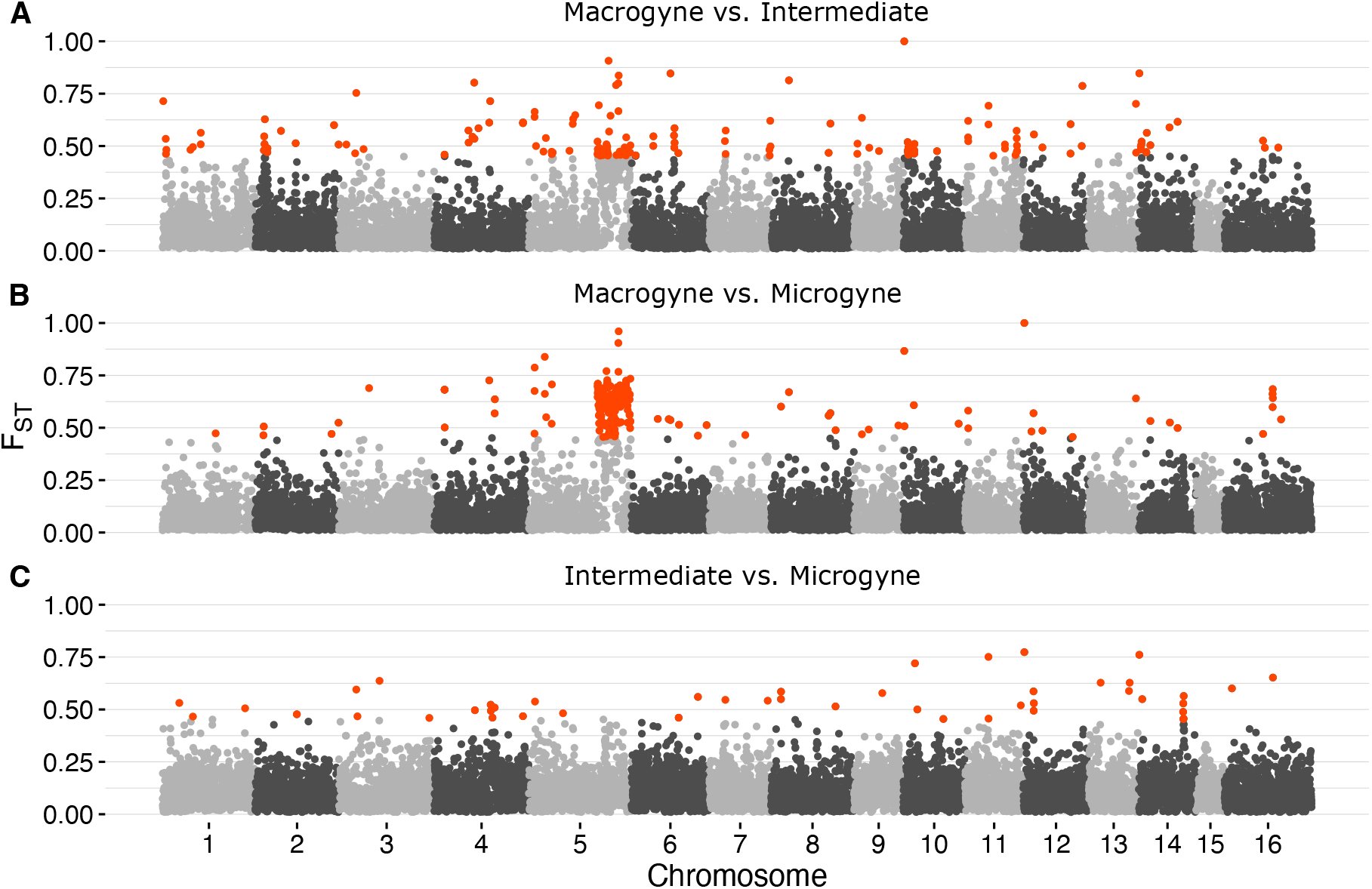
Genome-wide F_ST_ scans across 25 kb windows between the (**A**) macrogynes (n = 3; head width >1.08 mm) and intermediates (n = 4; head width 1.04-1.08), (**B**) macrogynes and microgynes (n = 10; head width <1.04), and (**C**) intermediates and microgynes. A 9 Mb region on chromosome 5 was significantly differentiated (values above the 95^th^ percentile are highlighted in red) between macrogynes and microgynes (**B**), somewhat differentiated between macrogynes and intermediates (**A**), but not between intermediates and microgynes (**C**).

We identified three distinct genotypes in the queen morph-differentiated region on chromosome 5, consistent with the expected signature of non-recombining supergene haplotypes, using a 50 kb rolling window PCA (Figure 2A). Among the queens/gynes in the intermediate PC1 cluster (Figure 2A; purple), most SNPs in the diverged region were heterozygous (60-63%), suggesting that these individuals carry one haplotype of each type. We refer to this group as “AB” genotypes. The two other PC1 clusters were less heterozygous (13-16%) and are interpreted to be homozygous for opposite supergene haplotypes and referred to as “AA” and “BB” (Figure 2; green and yellow). This grouping is supported by genotype counts of SNPs within this region, where the AA individuals were homozygous for the reference allele for most of the sites where all the AB individuals were heterozygous, while the BB individuals were mostly homozygous for the alternative allele (Figure 2B). Consistent with the expected patterns from non-recombining supergene haplotypes, the diverged region on chromosome 5 displayed elevated linkage disequilibrium compared to the rest of chromosome 5 and a genome-wide measurement (Figure 2C). The supergene polymorphism is also present among the 18 *M. ruginodis* workers, as shown by a rolling window PCA (n = 7 AA, n = 9 AB, n = 2 BB) (Figure S3; Table S1).

**Figure 2.**
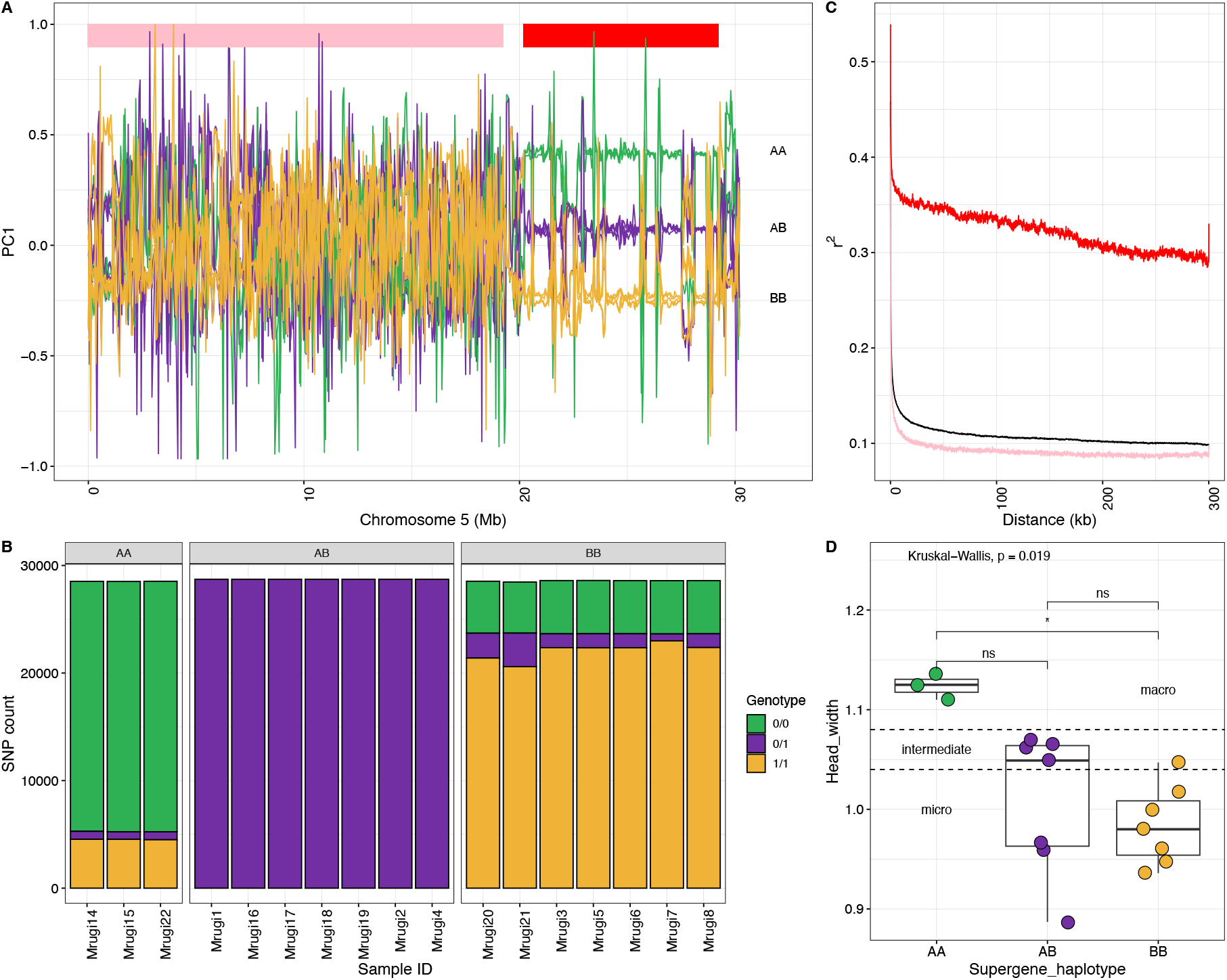
(**A**) Rolling window PCA1 values across chromosome 5 for all queens/gynes (n = 17), revealing three distinct supergene genotypes (“AA”, “AB”, and “BB”) on chromosome 5 between 20.2 and 29.2 Mb (region marked with a red rectangle). The line colour correspond to the supergene genotype of each queen/gyne (same as in **D**). (**B**) Genotype counts for SNPs in the supergene region that are heterozygous in all AB individuals (n = 53,817 SNPs). 0/0 = reference allele homozygous, 0/1 = heterozygous, 1/1 = alternative allele homozygous. (**C**) Decay of linkage disequilibrium within the chromosome 5 supergene region (red; region showed in **A**), the non-supergene region (pink; region showed in **A**) and a genome-wide estimate for all chromosomes (1-16) except chromosome 5 (black). (**D**) Head width measurements for each queen/gyne and supergene category. Data points are coloured according to the supergene genotype (see **A**).

Previous studies have shown that queen size is a genetically inherited trait in *M. ruginodis* (Wolf, 2016), and here we provide evidence of the genetic architecture underlying this phenotypic divergence. The three macrogynes all had the AA genotype, while the intermediates and microgynes had either AB or BB genotypes (Figure 2D; Table S1). We found a significant association between queen head width and supergene genotype (Kruskal-Wallis, H = 7.9173, df = 2, p-value = 0.019), with AA genotypes having a higher median head width (1.2 mm) than AB (1.05 mm) and BB genotypes (0.98 mm). After adjusting for multiple tests, only the difference between AA and BB genotypes was statically significant (Dunn test: z = 2.81; adj. p = 0.015; Figure 2D).

### PCR genotyping supports link between supergene and queen size

To verify the association between supergene genotype and queen size, we collected additional *M. ruginodis* queens (n = 78; Table S1) and determined their supergene genotype using a diagnostic PCR primer pair (see Methods). Of the 78 *M. ruginodis* queens, 11 were classified as AA homozygotes, 29 as AB heterozygotes, and 38 as BB homozygotes (Table S1; Figure S4-6). Again, we found statistical differences in head width between the three supergene genotypes (Kruskal-Wallis: H = 37.039, df = 2, p-value = <0.0001). The AA genotypes were significantly larger than both AB (Dunn test: z = 4.73; adj. p-value < 0.001) and BB (Dunn test: z = 6.08; adj. p-value < 0.001) genotypes (Figure 3). There was, as before (Figure 2D), no statistical difference between the head sizes of AB and BB genotypes (Dunn test: z = 1.65; adj. p-value = 0.10; Figure 3). In contrast to the analysis of only the whole-genome sequenced queens and gynes (Figure 2D, there was some overlap between the head width distribution of the AA genotypes and those of the two other groups (Figure 3). This may be the result of some error associated with the head width measurements, or that environmental factors and/or other genetic elements may contribute to *M. ruginodis* queen size. Similar partial overlaps in supergene-controlled queen size have been found in *Formica cinerea* (Scarparo et al., 2023).

**Figure 2.**
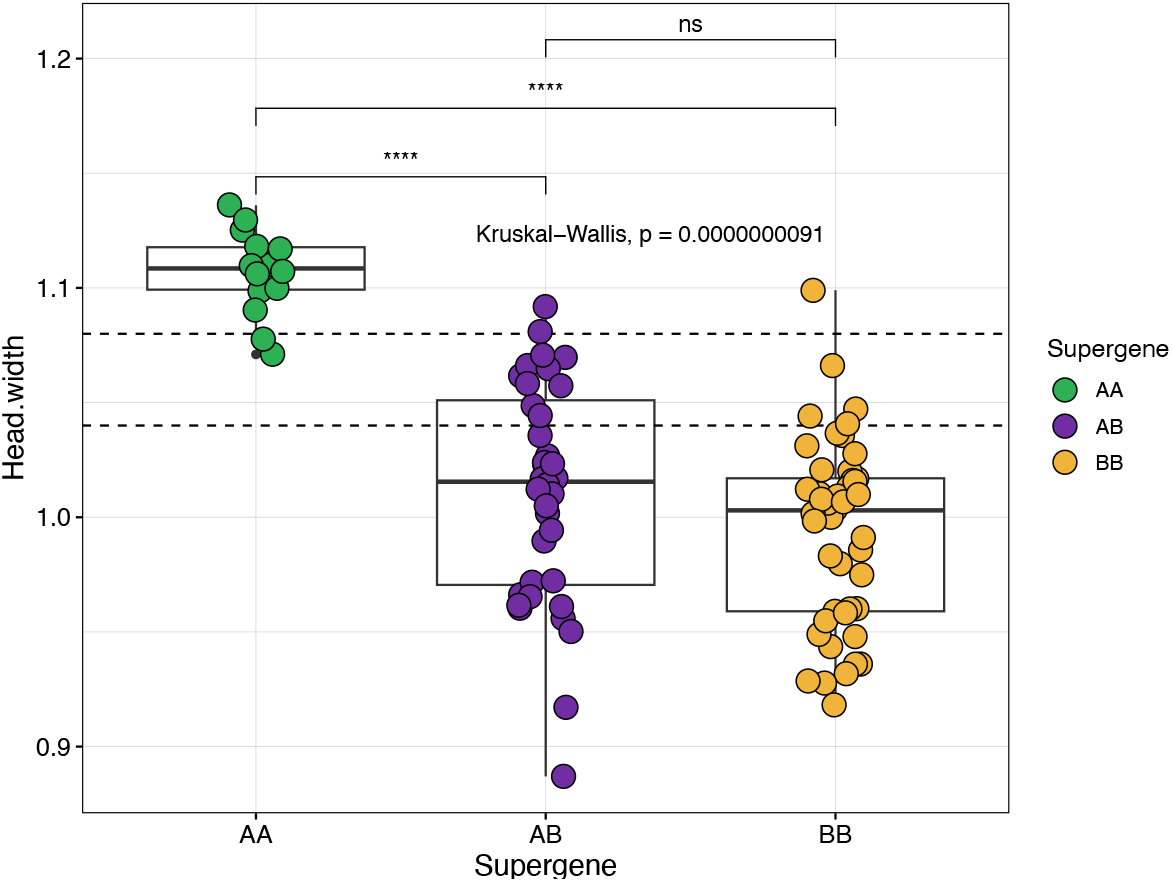
Head width and supergene genotype for the 95 queens/gynes (n = 17 WGS sequenced, n = 78 PCR genotyped). The dashed lines mark cut-off values between macrogynes, intermediates and microgynes (as in Figure 2D).

### The supergene determines social colony organization

Next, we investigated the association between supergene and colony queen number. The number of captured queens in the 31 colonies ranged between 1 and 25 (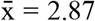Table S1; Figure 4), with 20 single-queen and 11 multiple-queen colonies (Figure 4A, B). Across the 20 single-queen colonies, 14 had AA queens while 6 had AB or BB queens (Figure 4A). Not a single AA genotype was found among the 69 queens from the multiple-queen colonies (Figure 4B). The genotypes in the multiple-queen colonies were either fully AB (n = 3), fully BB (n = 2) or a mixture (n = 6; Figure 4B). The association between the AA genotype and single-queen nesting was highly significant (Fisher’s exact test, p = 0.00015). In the single-queen colonies, the six non-AA queens were significantly smaller than the AA queens (Dunn test: z = 4.36; adj. p-value = <0.0001; Figure 4C), but of similar size to the non-AA queens in the multiple-queen colonies (Dunn test: z = 1.11; adj. p-value = 0.268; Figure 4C, D).

**Figure 4.**
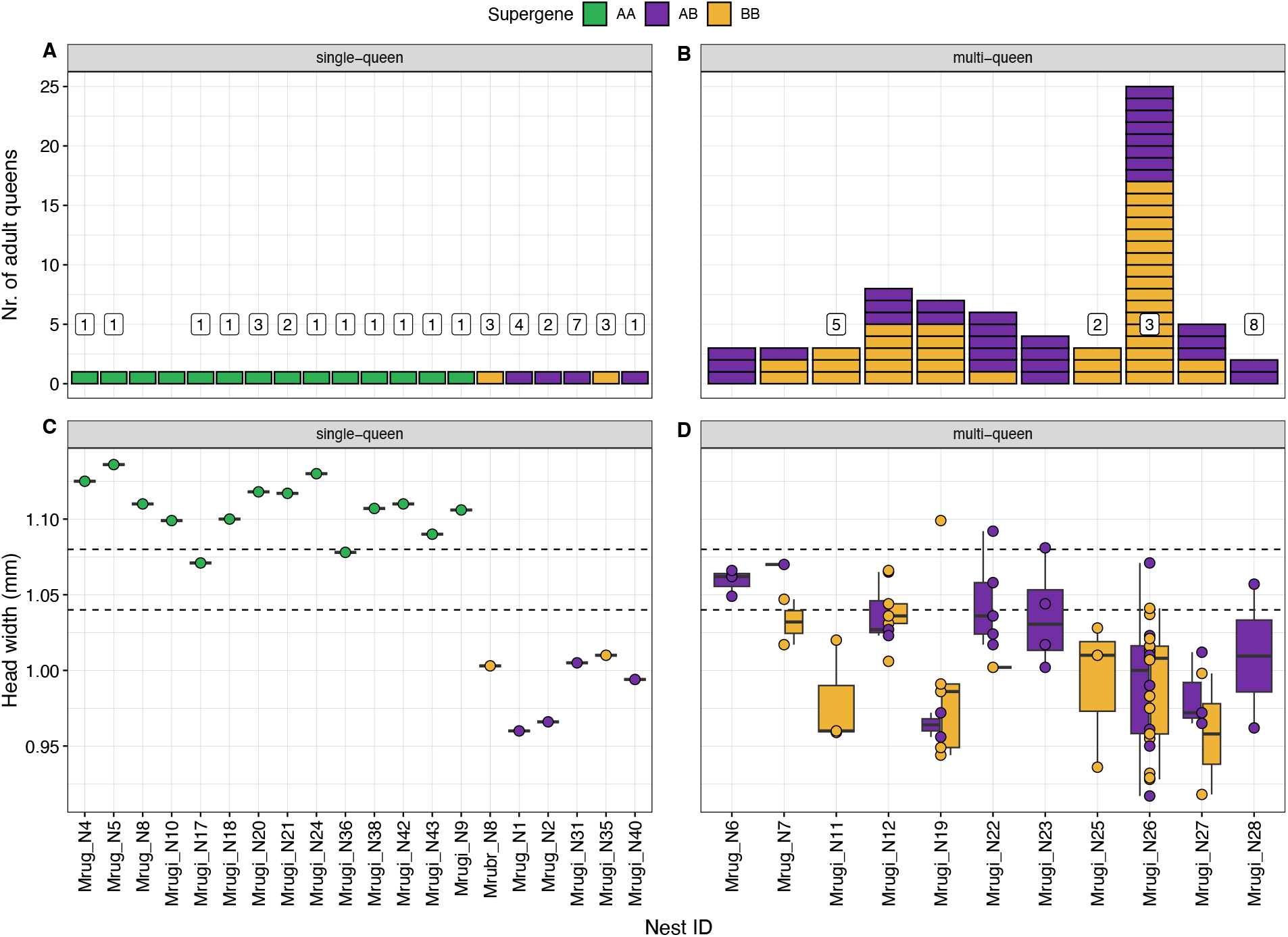
(**A,B**) Number of captured (wingless, i.e., adult) queens and (**C,D**) head width per colony (n = 31). The left-side panels (**A,C**) shows only colonies with a single queen (n = 20) and the right-side panels (**B,D**) show colonies where multiple queens were found (n = 11). The number within white boxes in panel **A** and **B** show the number of estimated matrilines based on worker microsatellite data. The dashed lines in **C** and **D** mark cut-off values between macrogynes, intermediates and microgynes (as in Figure 2D).

To further investigate the true sociogenetic structure of single-queen colonies, we clustered worker microsatellite genotypes to matrilines using the software Colony (see Methods). Most of the single-queen AA colonies had only one matriline (n = 10/12; Figure 4A; Table S16). However, multiple matrilines were detected in two AA colonies (n = 2/12; 2-3 matrilines; Figure 4A; Table S16). In contrast, multiple matrilines were detected in almost all non-AA single-queen colonies (n = 5/6; 2-7 matrilines; Figure 4A; Table S16). All four multiple-queen colonies analyzed as a control had multiple matrilines (n = 4/4; 2-8 matrilines; Figure 4B; Table S16). There was a significant association between a single matriline (i.e., monogyny) and queen AA genotype, and between multiple matrilines (i.e. polygyny) and AB or BB queen genotypes (Fisher’s exact test, p = 0.001905).

There are two potential explanations for the non-perfect association between supergene genotype and colony queen number. The first is that single-queen colonies can be headed by small non-AA queens. This seems unlikely, however, as smaller queens are typically less fecund (Brian, 1969; Keller, 1988; Vargo & Fletcher, 1989) and smaller colonies would be at disadvantage when competing with larger ones. Therefore, we argue that the single-queen colonies with non-AA genotypes are most likely polygynous, where additional queens may have died, moved, or escaped sampling. The second explanation is that single-queen colonies with multiple matrilines are actually polygynous. This could be a result of *serial polygyny*, recruitment of a replacement queen when the resident queen dies (Elmes, 1991; Seppä, 1994), with a transitory phase where offspring of multiple queens coexist. In *Myrmica*, this likely occurs as a result of the unusually long developmental time of sexual offspring (1 year) (Brian, 1983) combined with a relatively short queen life span (∼1.5 years) (Elmes, 1980; Evans, 1995; Seppä, 1994), which often leaves colonies queenless and prone to recruit new, even unrelated, queens (Evans, 1996).

One potential issue is that the mutation rate in microsatellite markers is high, and they are prone to genotyping errors. Even though this can be controlled for in the Colony analysis, it may affect sibship assignments even at low frequencies (J. Wang, 2018). It should also be noted that the result of the Colony analysis was partly ambiguous and the statistical support for finding multiple matrilines in single-queen colonies was not always strong (Table S16). In fact, worker genotypes in both single-queen colonies with multiple matrilines detected could also be explained with a single multiply mated queen (Table S10). Future studies using a larger sample of workers and genome sequencing data will reveal the relationship between true monogyny and the AA genotype with higher accuracy.

### Candidate genes for queen size and/or colony social organization

Supergenes are characterized by low recombination between haplotypes, meaning that genes within these regions are inherited as a single unit. This tight linkage makes it difficult to determine which specific genes contribute to the trait controlled by the supergene. Genes with many haplotype-specific SNPs—variants that are fixed for opposite homozygous alleles in AA and BB individuals, while heterozygous in AB individuals—are more likely than others to play a role in the trait associated with the supergene. To narrow down potential candidate genes, we counted the number of haplotype-specific SNPs within coding sequence (CDS) regions for each gene within the 9 Mb supergene region (Table S17). The two genes with the most haplotype-specific SNPs were *ABCA13* (“ATP-binding cassette, sub-family A”; n = 57) and *Twitchin* (n = 45; Table S17). In *Drosophila*, knockdown of *ABCA13* has been shown to affect social behavior, leading to an increased distance between individuals (Ueoka et al., 2018). *Twitchin* plays a regulatory role in muscle contraction, and according to one study insect flight muscles (Ayme-Southgate et al., 1991). Interestingly, in two obligately polygynous species of the *Formica rufa* group, one of the few remaining genes from a polygyny-associated supergene is *Zasp52*, a flight-muscle gene (Liao et al., 2016; Sigeman et al., 2024). Given that dispersal strategy is a key factor in alternative reproductive strategies in ants, differences in genes affecting flight capability would not be unexpected. However, additional data is needed to confirm the specific roles of these genes in shaping dispersal and social structure.

### Minimal shared synteny with other ant social supergenes

Social supergenes controlling queen number have now been found in six lineages of ants; *Cataglyphis, Formica, Leptothorax, Myrmica, Pogonomyrmex*, and *Solenopsis* (Braim, 2015; Errbii et al., 2024; Lajmi et al., 2024; Purcell et al., 2014; J. Wang et al., 2013) (this study). Previous studies have reported no gene overlap between the supergenes of *Formica, Solenopsis*, and *Pogonomyrmex* (Brelsford et al., 2020; Errbii et al., 2024), suggesting that they arose through independent evolutionary processes from different starting chromosomes. A recent study showed that the supergene of *Cataglyphis* shares synteny with the *Solenopsis* supergene (Lajmi et al., 2024). However, since *Cataglyphis* is more closely related to *Formica* than *Solenopsis*, the authors proposed that the supergene arose independently of the one in *Solenopsis* (Lajmi et al., 2024). For the last supergene, detected in *Leptothorax* (Braim, 2015), the size and gene content are not yet well characterized (Kay et al., 2022).

Synteny analyses of the *Myrmica* supergene against those of *Solenopsis, Formica*, and *Pogonomyrmex* reveal very little gene overlap (Figure 5). Only 12 *Myrmica* genes are shared between any of the other supergene regions, between *Myrmica* and *Formica* (Figure 5B; Table S18). Three of these shared genes had haplotype-specific SNPs within their CDS regions, although few: Mrub_g18249_i1 (n = 1), Mrub_g05933_i1 (n = 1), and Mrub_g08367_i1 (n = 3; Table S18). This suggests that the *Myrmica* supergene evolved independently and from the others. Note that the *M. rubra* reference genome was used in this analysis as there is no *M. ruginodis* reference genome available. The gene order within the *M. ruginodis* supergene region may therefore differ from the one shown here. Future studies using a high-quality reference genome from *M. ruginodis* will better characterize the structure and gene order of the supergene.

**Figure 5.**
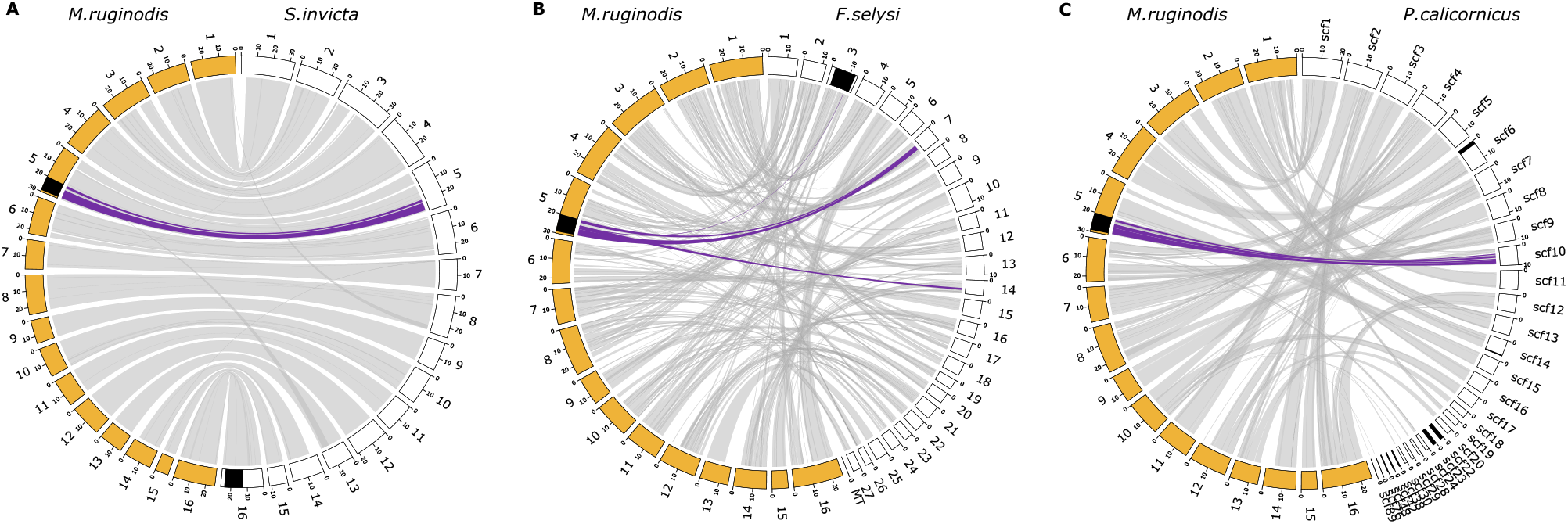
Synteny plots between M. ruginodis and (**A**) S. invicta, (**B**) F. selysi, and (**C**) P. californicus. M. ruginodis chromosomes are in yellow, and in white for the comparison species. Social supergene regions in each species are highlighted in black. Gray links show genome-wide syntenic regions between reference genomes. Purple links highlight syntenic gene pairs within 10kb genome windows in the M. ruginodis reference genome containing haplotype-specific SNPs. There is no gene overlap between the supergene regions of M. ruginodis and (**A**) S. invicta, or (**C**) P. californicus. Between M. ruginodis and (**B**) F. selysi, however, a small fraction of genes (n = 12; Table S18) are shared between the supergene regions. There is, however, no gene content shared between the Myrmica supergene and F. selysi chromosome 9, which is homologous to the chromosome carrying the queen-size controlling supergene in polygynous F. cinerea colonies (Scarparo et al., 2023). The C. niger supergene shares synteny with the S. invicta supergene region on chromosome 16 (Lajmi et al., 2024), but no precise supergene genome coordinates are available for the currently published reference genomes of this species.

## Conclusion

We demonstrate that a 9 Mb supergene is associated with the queen morphs and colony social organization in *M. ruginodis*, so that one genotype of the supergene (AA) is found exclusively in large queens in monogynous colonies, and two others (AB, BB) in smaller queens in polygynous colonies. This suggests that colony social organization and queen morph are genetically controlled by this supergene in *M. ruginodis*. Through comparative genomics, we show that the novel supergene does not share synteny with other ant supergenes, except for 12 genes that are shared between the *Myrmica* and *Formica* supergenes. The high frequency of all supergene genotypes (AA, AB, and BB) suggests that no combination carries severe fitness consequences. Our findings align with the patterns found in previous research on the queen-size dimorphic ant *M. ruginodis*, while providing more accurate information on the genetic mechanism maintaining distinct social organization in the two morphs.

## Notes

### Competing Interest Statement

The authors have declared no competing interest.

### Summary of Updates

This version contains a revision of Figure 1.

