## Supplementary Figures for "A novel supergene controls queen size and colony social organization in the ant *Myrmica ruginodis*"

### Supplementary Figures S1-S6

#### A novel supergene controls queen size and colony social organization in the ant *Myrmica ruginodis*

Hanna Sigeman[1,2], Perttu Seppä[3], Philip A. Downing[1], Matthew Webster[2], Heikki Helanterä[1,4], and Lumi Viljakainen[1]

[1]Ecology and Genetics Research Unit, University of Oulu, Finland
[2]Department of Medical Biochemistry and Microbiology, Uppsala University, Sweden
[3]Organismal and Evolutionary Biology Research Programme, University of Helsinki, Finland
[4]Tvärminne Zoological Station, University of Helsinki, Finland

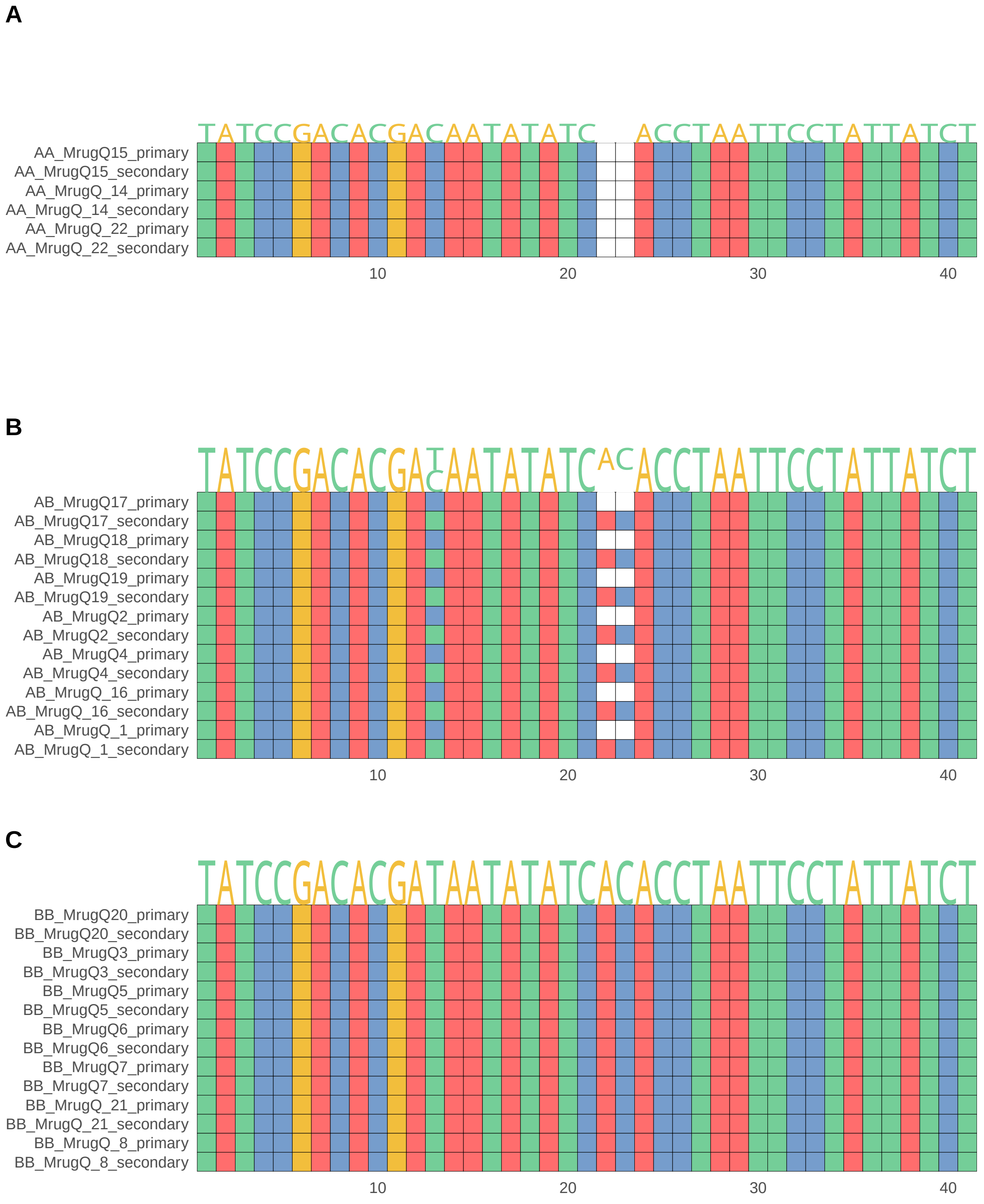


Figure S1: DNA motifs from the supergene haplotype diagnostic primer pair IL16 for 17 whole-genome sequenced queens/gynes. For each individual, electropherogram data from Sanger sequencing were phased into a 'primary' and 'secondary' sequence. The supergene genotype can be inferred through a B-specific 2bp insertion (position 22-23). The queens/gynes are grouped according to their supergene genotype: (**A**) AA, (**B**) AB,(**C**) BB (Table S1).

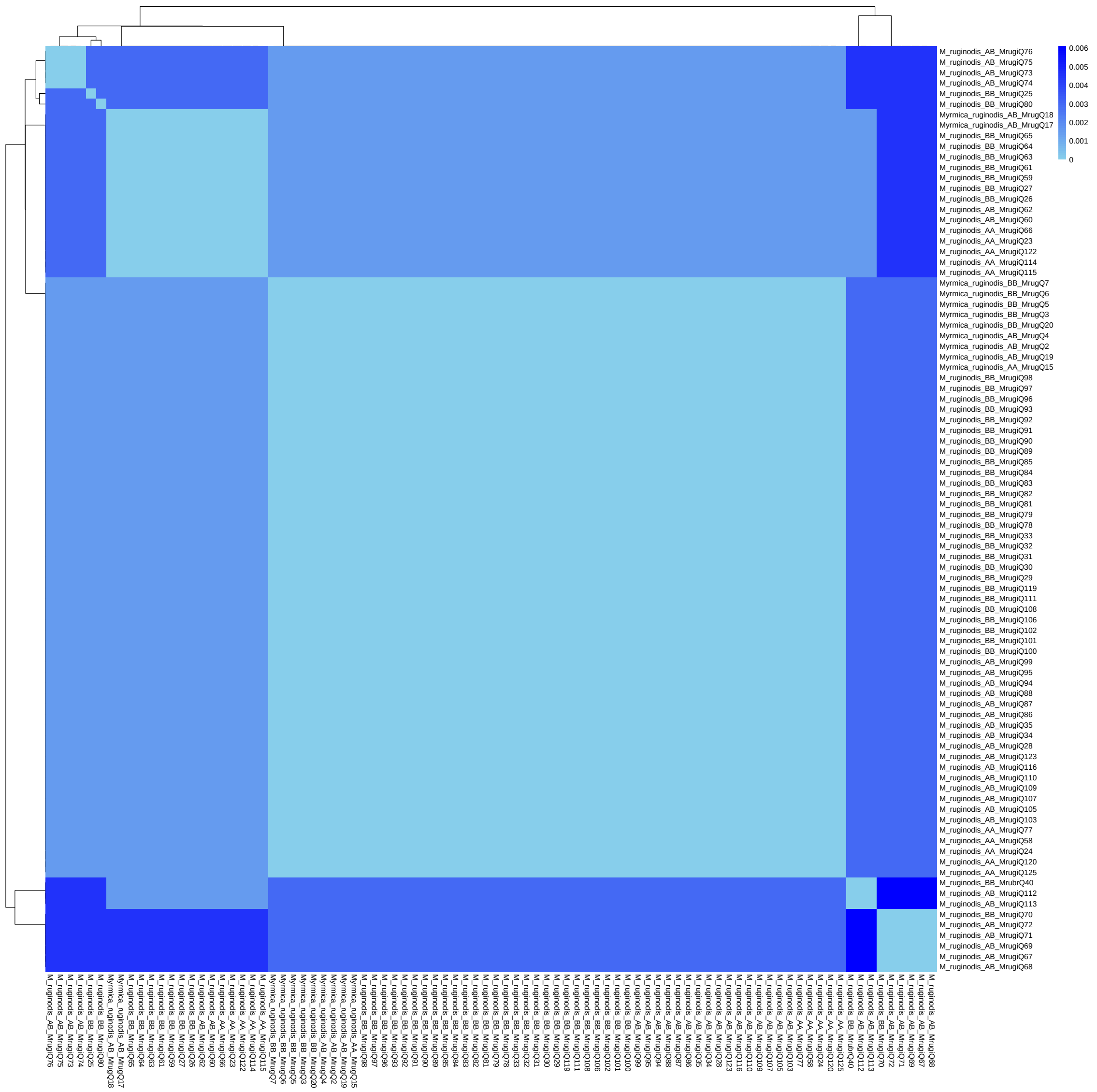


Figure S2: Heatmap showing pairwise sequence distances between COI sequences from *M. ruginodis* queens/gynes. The sequence distances were small between all individuals, ranging between 0 and 0.006, confirming that all individuals belong to the same species.

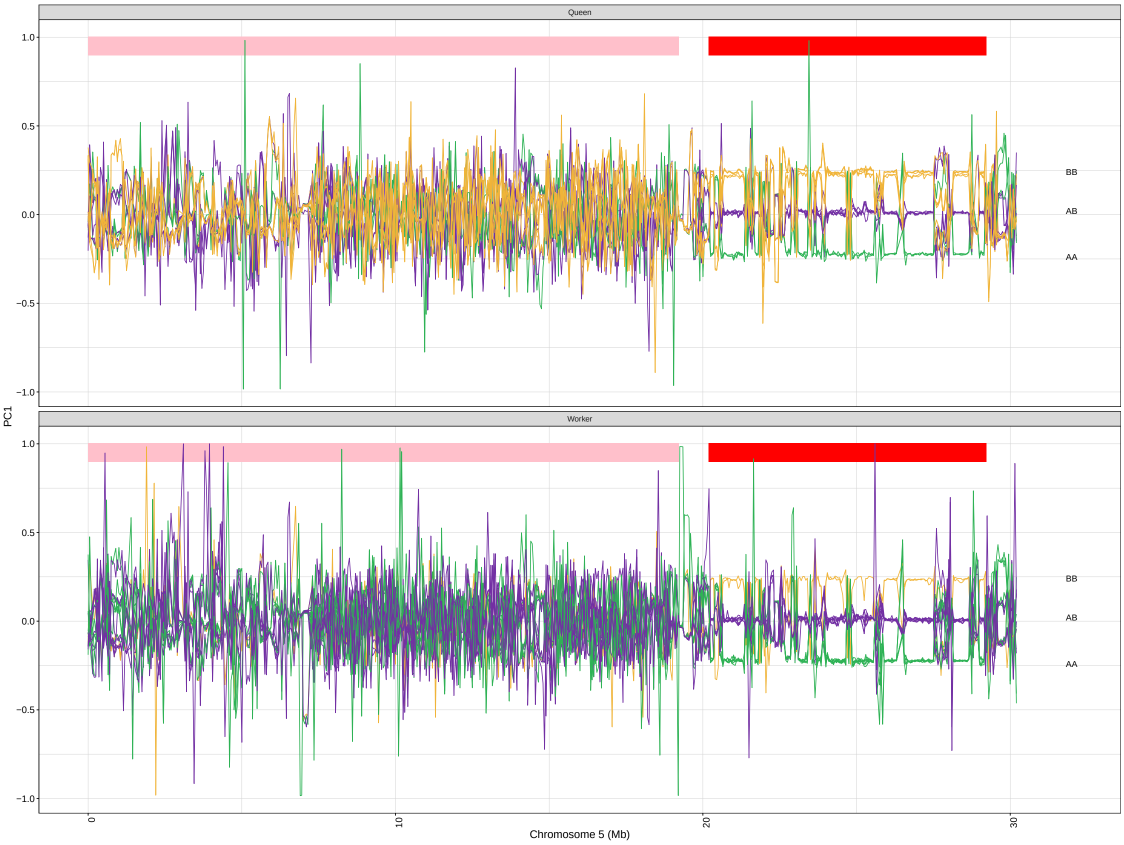


Figure S3: Rolling PCA plot across chromosome 5, showing the three supergene genotypes AA, AB, and BB. The upper panel show data for the 17 whole-genome sequenced *M. ruginodis* queens/gynes (same individuals as Figure 2A) and the bottom panel show data from 18 *M. ruginodis* workers. See Table S1 for sample details.

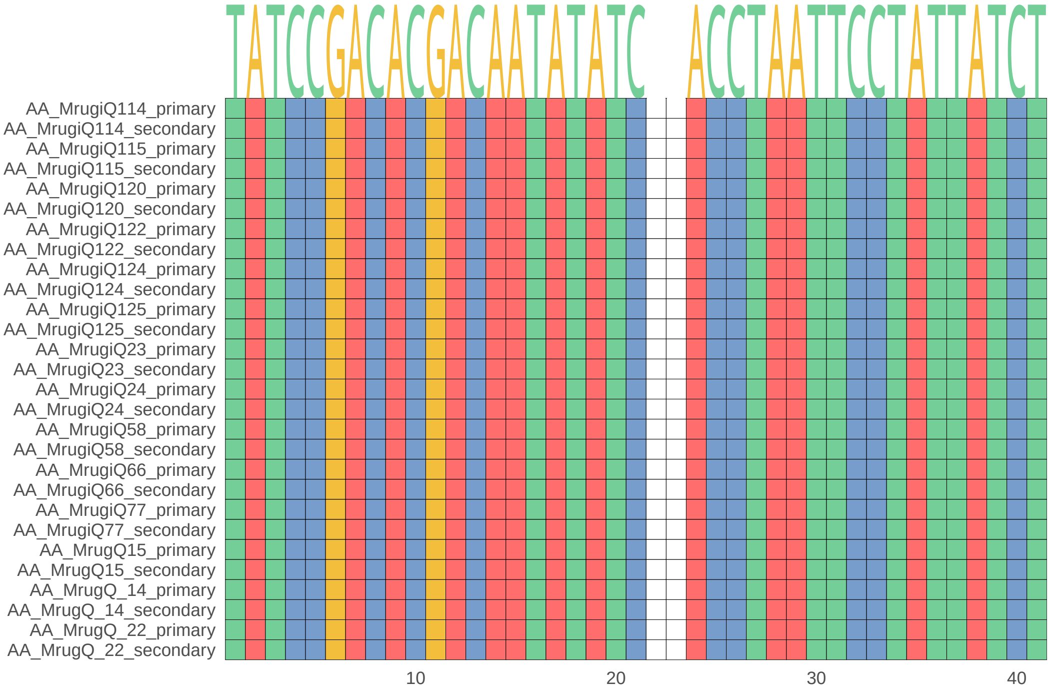


Figure S4: All additional (non-whole genome sequenced) AA queens genotyped using the IL16 supergene diagnostic PCR primers. See Table S1 for sample details.

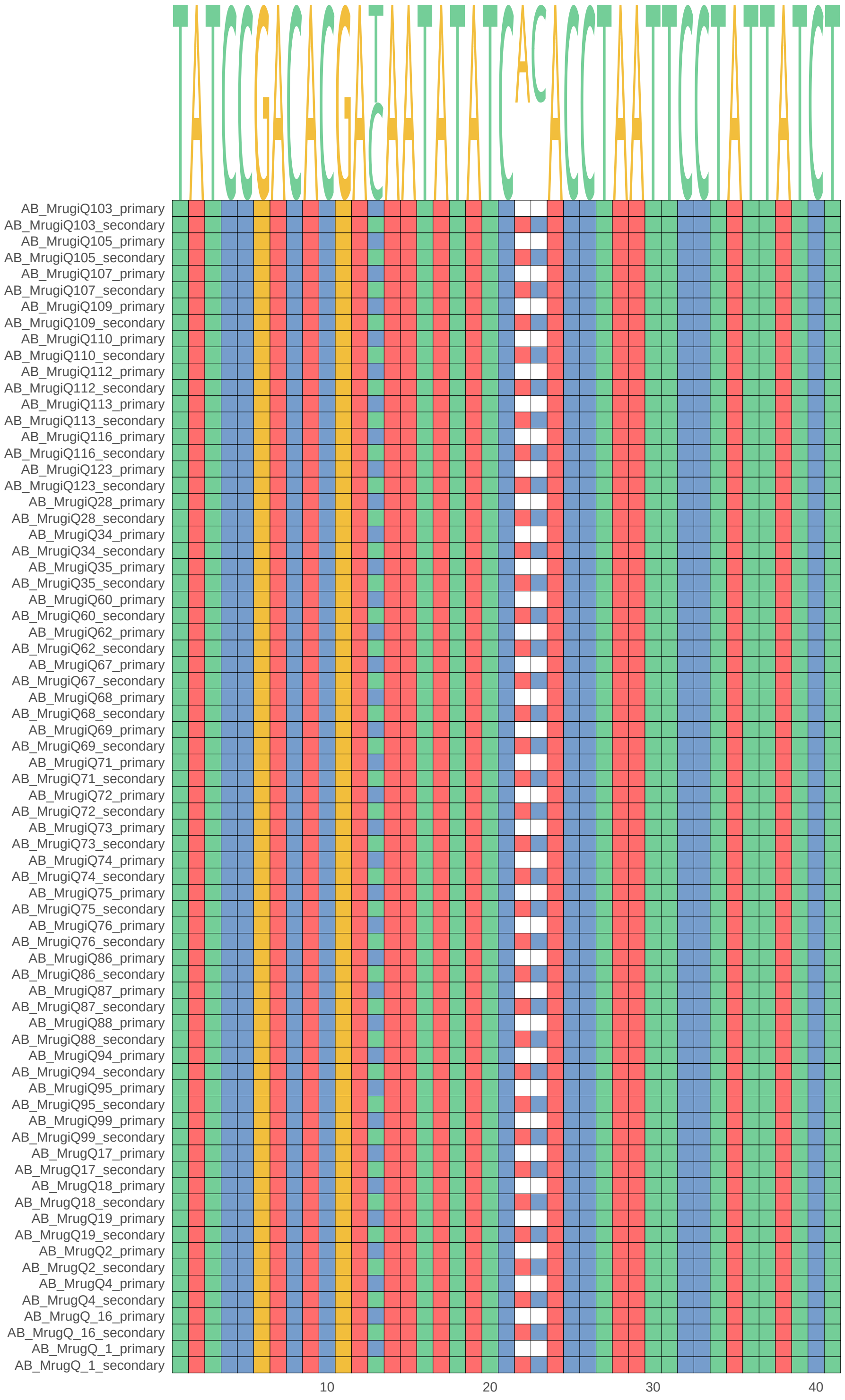


Figure S5: All additional (non-whole genome sequenced) AB queens genotyped using the IL16 supergene diagnostic PCR primers. See Table S1 for sample details.

C

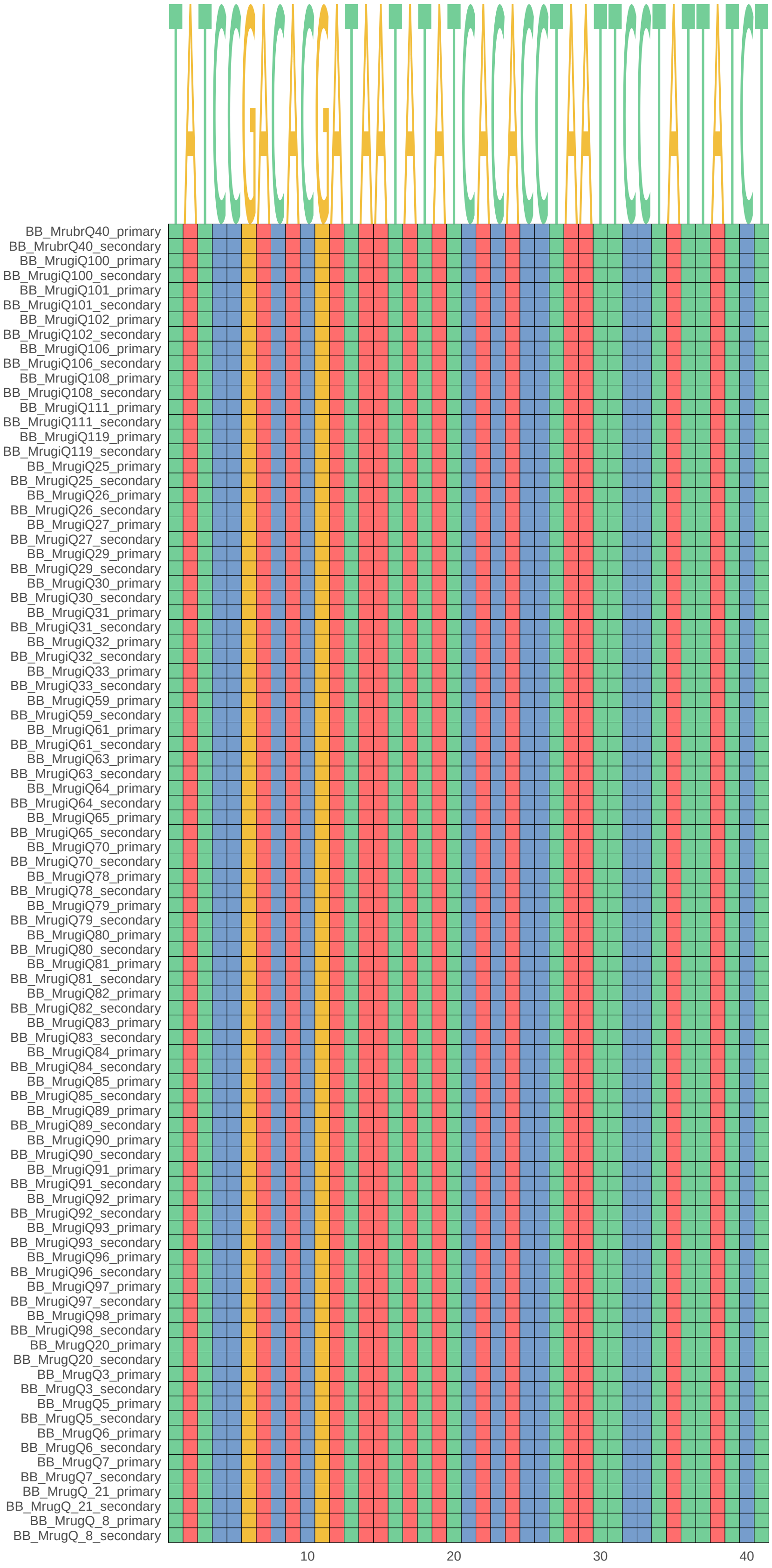


Figure S6: All additional (non-whole genome sequenced) BB queens genotyped using the IL16 supergene diagnostic PCR primers. See Table S1 for sample details.
